# Development of 5-FU-modified tumor suppressor microRNAs as a platform for novel microRNA-based cancer therapeutics

**DOI:** 10.1101/2022.06.23.497408

**Authors:** John G. Yuen, Andrew Fesler, Ga-Ram Hwang, Lan-Bo Chen, Jingfang Ju

## Abstract

MicroRNA (miRNAs) are pleiotropic post-transcriptional modulators of gene expression. Its inherently pleiotropic nature make miRNAs strong candidates for the development of cancer therapeutics, yet despite its potential, there remains a challenge to deliver nucleic acid-based therapies into cancer cells. We developed a novel approach to modify miRNAs by replacing the uracil bases with 5-fluorouracil (5-FU) in the guide strand of tumor suppressor miRNAs, thereby combining the therapeutic effect of 5-FU with tumor suppressive effect of miRNAs to create a potent, multi-targeted therapeutic molecule without altering its native RNA interference (RNAi) function. To demonstrate the general applicability of this approach to other tumor suppressive miRNAs, we screened a panel of 12 novel miRNA mimetics in several cancer types including leukemia, breast, gastric, lung, and pancreatic cancer. Our results show that 5-FU-modified miRNA mimetics have increased potency (low nM range) in inhibiting cancer cell proliferation and that these mimetics can be delivered into cancer cells without delivery vehicle both *in vitro* and *in vivo*, thus representing significant advancements in the development of therapeutic miRNAs for cancer. This work demonstrates the potential of fluoropyrimidine modifications that can be broadly applicable and may serve as a platform technology for future miRNA and nucleic acid-based therapeutics.

## Introduction

MicroRNAs (miRNAs) are a class of short non-coding RNA that were discovered in 1993 and were observed to have critical regulatory functions, regulating protein biosynthesis via direct interaction with the 3’ untranslated region (3’-UTR) of target mRNA transcripts.^1,2^ MiRNA are key mediators of the RNA interference (RNAi) pathway and its interactions are largely governed by its seed sequence on the 5’ end of the miRNA. They are thus pleiotropic regulators of gene expression, where one particular miRNA can interact with multiple mRNA target transcripts. There are numerous studies demonstrating the critical roles of miRNAs in human disease and miRNA expression is often dysregulated in cancer. Due to their impact on numerous biochemical pathways, different miRNAs have been observed to either promote or inhibit tumorigenesis and tumor growth in cell context-dependent manner. These miRNAs are referred to as oncogenic miRNAs (oncomiRs) and tumor suppressor miRNAs respectively.^3^ miR-15 was the first miRNA to be found either absent or downregulated in cancer and it was subsequently discovered that it modulates apoptosis by directly regulating BCL2 expression.^4,5^ Since then, numerous cancer types have been shown to have dysregulated tumor suppressor miRNA expression. The functional significance of miRNAs in cancer has also been further studied, and tumor suppressor miRNAs that regulate key oncogenic pathways—such as apoptosis, proliferation, autophagy, cell cycle, and epithelial-to-mesenchymal (EMT) transition—have been identified and studied.^6-13^ Experimental evidence that demonstrates the key role of miRNAs in these pathways reveals the therapeutic potential for tumor suppressor miRNAs.

There are, however, several hurdles need to be overcome to realize the therapeutic potential of miRNAs. One of the major bottlenecks in the development of nucleic acid-based medicine is the efficient intracellular delivery of these molecules. Tremendous effort in the past several decades has been devoted to developing delivery vehicle technologies and various lipid-based nanoparticles—including liposomes, micelles, and dendrimers—have grown in popularity.^14,15^ Many of these lipid nanoparticles (LNPs) function as cationic polymers that facilitate the transport of oligonucleotides across the plasma membrane. Despite the advancements in lipid-based delivery systems, there are still barriers to its use. LNPs still face several challenges such as their instability, rapid systemic clearance, and toxicities—including their induction of immunostimulatory responses. In preclinical studies, LNPs are known to exhibit some cytotoxicity in various cell lines and mice treated with some formulations of positively-charged LNPs showed increased liver enzymes and body weight loss.^16,17^ In humans, anti-polyethylene glycol (PEG) antibodies are generated from pegylated drugs, including LNP formulations, and PEG-induced complement activation has also been observed.^18-21^ Toll-like receptors (TLRs) are involved in the innate immune response and typically recognize molecular patterns associated with pathogens.^22^ TLR-4 activation has also been reported with the use LNPs.^16^ These toxicities are particularly problematic as they are often observed when LNPs are delivered systemically, the main route of delivery in cancer therapy.^23-25^

Other approaches to enhance delivery include the modification of the nucleic acids themselves. Modifications of the sugar-phosphate backbone and/or the base have been shown to enhance delivery, stability, and potency of various nucleic acid-based therapeutics.^26^ 5-fluorouracil (5-FU) is a pyrimidine analog and is converted to fluorodeoxyuridylate (FdUMP) to inhibit the sole *de novo* biosynthesis of thymidylate by the formation of suicide ternary complex with its target enzyme protein thymidylate synthase (TS) along with tetrahydrofolate.^27^ Due to this antimetabolite effect, 5-FU has been used as a major chemotherapeutic agent for many cancer types. 5-FU can be incorporated into the miRNA molecule and we have previously taken this approach with three tumor suppressor miRNAs—miR-15a, miR-129, and miR-489—and have demonstrated that 5-FU modification of these miRNAs are an effective and potent cancer therapeutic in *in vivo* mouse models of colon, pancreatic, and breast cancer.^28-31^ Additionally, this modification confers increased intracellular stability of the miRNA, a novel ability to be delivered into cells without use of delivery vehicle, and retention of target specificity. This modification strategy greatly overcomes some of the bottlenecks of nucleic acid-based drug development by improving the deliverability of miRNAs as well enhancing its potency. To demonstrate the general applicability of this concept, we took this unique 5-FU modification-approach to develop a miRNA strategy that can be implemented as a platform for miRNA-based cancer therapeutics.

In this study, we selected 12 well-studied tumor suppressor miRNAs (hsa-let-7a, hsa-miR-15a, hsa-miR-34, hsa-miR-129, hsa-miR-140, hsa-miR-145, hsa-miR-194, hsa-miR-200a, hsa-miR-200b, hsa-miR-200c, hsa-miR-215, and hsa-miR-506) as candidates to apply this modification strategy.^6-13,32-37^ These tumor suppressor miRNAs are often dysregulated in cancer cells and play important roles in multiple pathways regulating the cell cycle, apoptosis, EMT, metastasis, and drug resistance.^38^ We substituted the uracil bases on the guide strand of the miRNAs with 5-FU and screened these 5-FU-modified miRNA mimetics for efficacy in various cancer types. The passenger strand of miRNA was left unmodified to avoid potential off-target effects and to preserve the function of miRNA. Notably, 5-FU modification does not affect Watson-Crick base-pairing, as the fluorine substituted for the hydrogen at the 5-carbon is not involved in normal hydrogen bonding between nucleobases. This unique strategy combines the therapeutic effect of 5-FU with the tumor suppressor function of miRNAs to inhibit multiple oncogenic targets and pathways of cancer cells. Our results show that these 5-FU-modified miRNA mimetics were able to reduce cancer cell proliferation with enhanced potency while preserving target specificity. Lastly, the therapeutic effects were achieved without the aid of any delivery vehicle both *in vitro* and *in vivo*.

## Results

### Sequence and structure of the miRNA mimetics with 5-FU modification

To evaluate 5-FU modification as a universal strategy for all tumor suppressor miRNAs and for multiple major cancer types, we selected twelve tumor suppressor miRNAs that are well studied in several cancer types, including in gastric cancer, lung cancer, breast cancer, leukemia, and pancreatic cancer: hsa-let-7a, hsa-miR-15a, hsa-miR-34, hsa-miR-129, hsa-miR-140, hsa-miR-145, hsa-miR-194, hsa-miR-125, hsa-miR-200a, hsa-miR-200b, hsa-miR-200c, and hsa-miR-506 (herein after referred to without their hsa-prefix which refers to their human species of origin). We designed a panel of 5-FU-modified miRNAs by substituting the uracil bases of the guide strand of the mature miRNA with the antimetabolite nucleoside analog 5-FU **(Figure 1)**.

**Figure 1.**
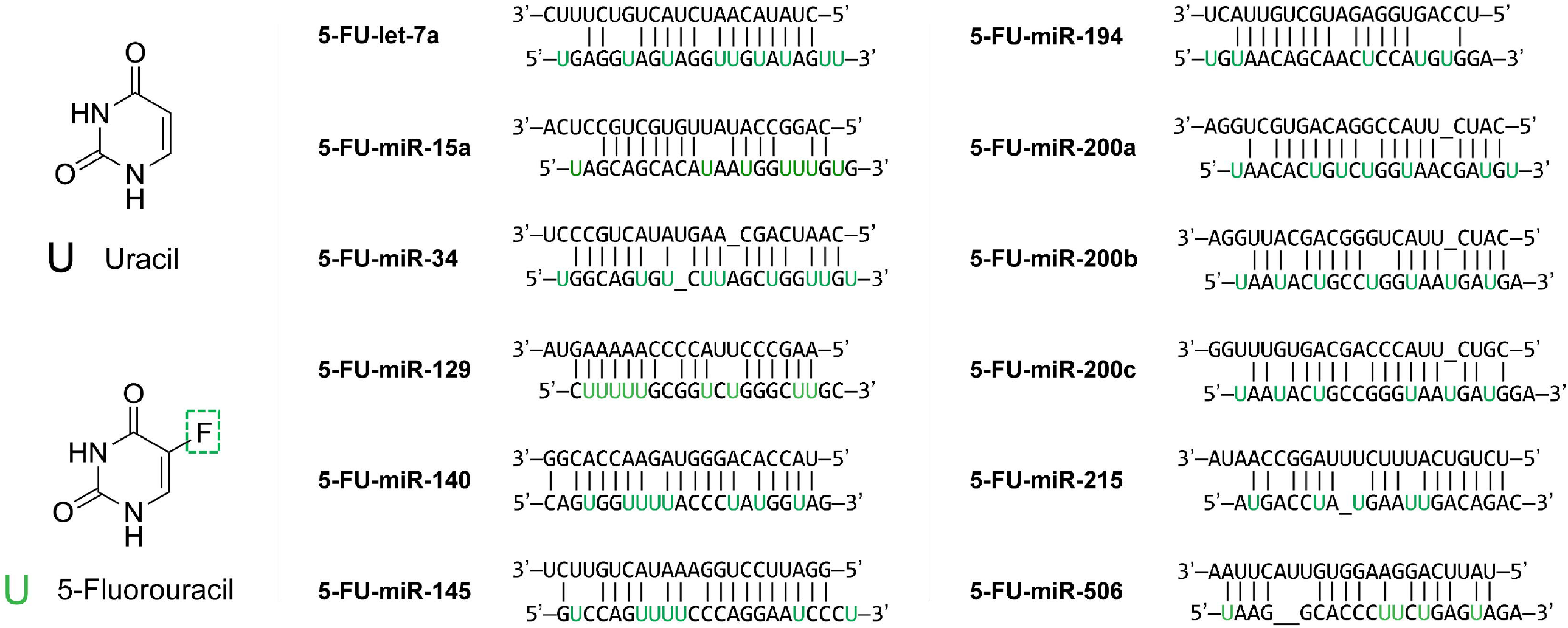
Panel of 5-fluorouracil-modified miRNAs. The sequences of twelve tumor suppressor miRNAs—let-7a, miR-15a, miR-34, miR-129, miR-140, miR-145, miR-194, miR-125, miR-200a/b/c, and miR-506—with their uracil bases of the 5’ strand of the mature miRNA substituted with the antimetabolite nucleoside analog 5-fluorouracil (5-FU).

### Toxicity of Nucleic Acid Delivery Vehicle

To demonstrate the cytotoxicity of delivery vehicles, both gastric cancer AGS and lung cancer A549 cells were treated with a polyethyleneimine-based lipid nanoparticle (*in vivo*-jetPEI®, Polyplus) and a dose-dependent increase in apoptosis was observed **(Figure 2)**. It is clearly evidenced that while PEI has minimal impact on cell death at low concentrations, PEI is cytotoxic at higher concentration and triggers apoptosis.

**Figure 2.**
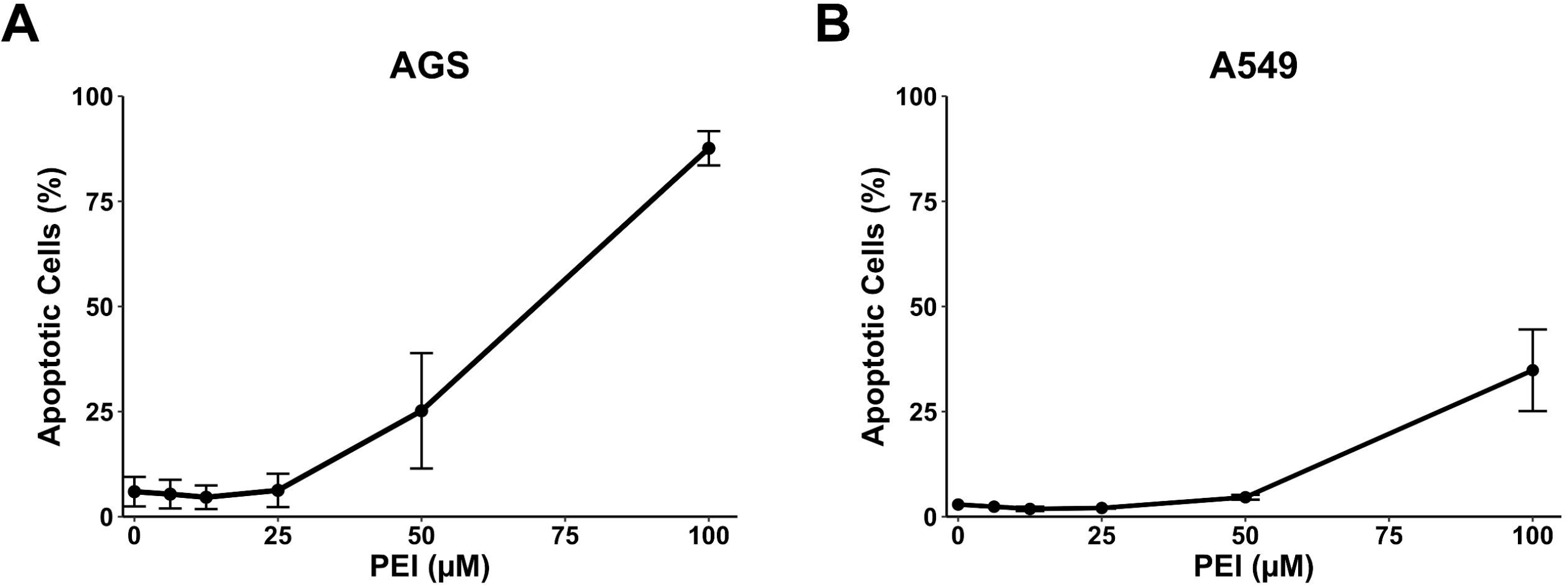
Nucleic acid delivery vehicles are inherently toxic. Common cationic lipid-based nucleic acid delivery vehicles such as polyethyleneimine (PEI) are toxic to cells. **(A)** AGS and **(B)** A549 cells exhibit dose-dependent apoptosis in the presence of PEI.

### 5-FU-modified miRNA mimetics display enhanced efficacy at inhibiting cancer proliferation and can enter the cell without the use of delivery vehicle

To systematically confirm that vehicle-free delivery of the 5-FU-modified miRNA mimetics is a reproducible, sequence-independent feature and that is also not dependent on the number of 5-FU substitutions, we explored the efficacy of our panel of 5-FU-modified miRNAs in different tumor types, we performed a screen by treating cells with 50 nM of the 5-FU-modified miRNAs without the use of delivery vehicle in the following cell lines: AGS gastric cancer cells, A549 lung cancer cells, SBKR3 breast cancer cells, REH leukemia cells, and AsPC-1, Hs766T, and PANC-1 pancreatic cancer cells **(Figure 3)**. Our results clearly show that while unmodified, control miRNAs have no effect due to the inability to cross the cell membrane, the 5-FU modified miRNA mimetics are all showing efficacy to inhibit cancer cell proliferation **(Figure 3)**.

**Figure 3.**
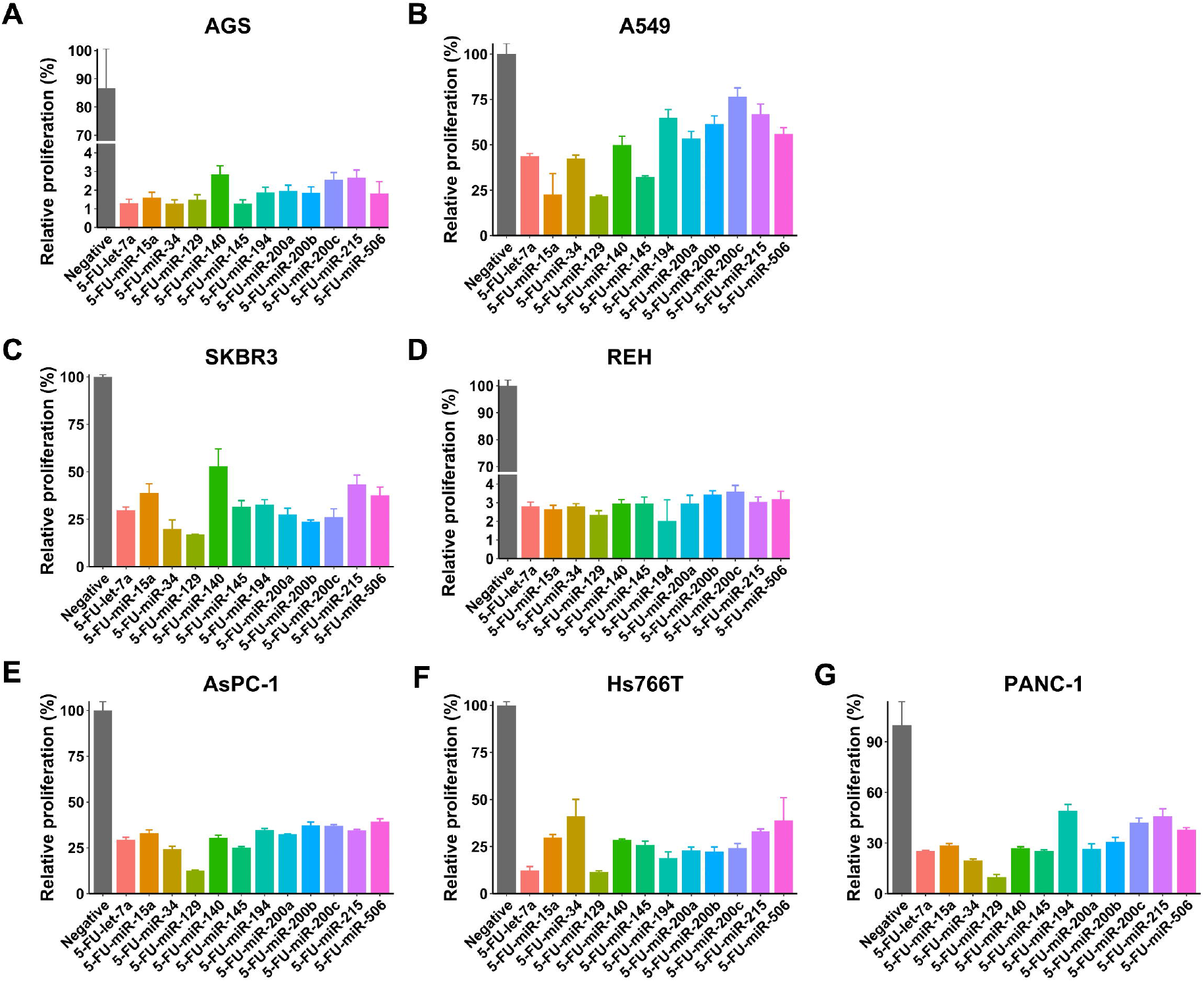
Vehicle-free treatment of 5-FU-modified miRNAs exhibit potent inhibition of tumor cell proliferation. A panel of twelve 5-FU-modified miRNA mimetics were screened for their effectiveness at inhibiting cell viability compared to negative control (scramble) miRNA at 50 nM without delivery vehicle. Several types of cancer cell lines were screened: **(A)** AGS gastric cancer cells, **(B)** A549 lung cancer cells, **(C)** SBKR3 breast cancer cells, **(D)** REH leukemia cells, and **(E)** AsPC-1, **(F)** Hs766T, and **(G)** PANC-1 pancreatic cancer cells.

In addition to vehicle free delivery, there is enhanced therapeutic efficacy of 5-FU-modified miRNA mimetics compared to their unmodified, native counterparts in both AGS cells **(Table 1)** and A549 cells **(Table 2)**. In AGS cells, there is an 8.4-fold increase in the inhibition of cell proliferation after 5-FU modification of let-7a (let-7a IC_50_ = 59.5 nM vs. 5-FU-let-7a IC_50_ = 7.1 nM) **(Figure 4A)** and a 4.3-fold increase for miR-145 (miR-145 IC_50_ = 20.9 nM vs. 5-FU-miR-145 IC_50_ = 4.9 nM) **(Figure 4B)**. Similarly, in A549 cells, there is a 6.3-fold increase in the inhibition of cell proliferation after modification of let-7a with 5-FU (let-7a IC_50_ = 143.5 nM vs. 5-FU-let-7a IC_50_ = 22.8 nM) **(Figure 4C)** and a 13.8-fold increase for miR-145 (miR-145 IC_50_ = 51.7 nM vs. 5-FU-miR-145 IC_50_ = 11.0 nM) **(Figure 4D)**.

**Table 1.**
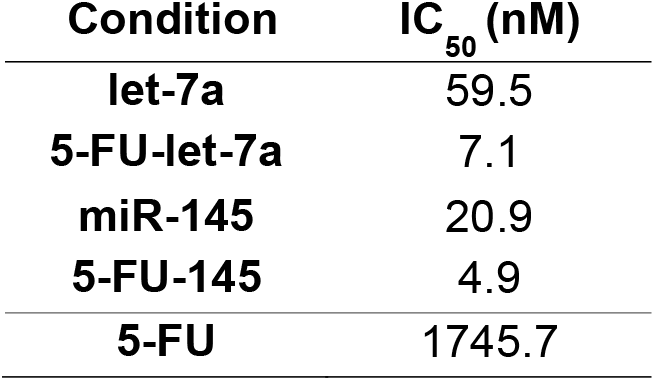
IC_50_ values of miRNAs in AGS cells.

| Condition | IC<sub>50</sub> (nM) |
| --- | --- |
| let-7a | 59.5 |
| 5-FU-let-7a | 7.1 |
| miR-145 | 20.9 |
| 5-FU-145 | 4.9 |
| 5-FU | 1745.7 |

**Table 2.** IC_50_ values of miRNAs in A549 cells.

| Condition | IC<sub>50</sub> (nM) |
| --- | --- |
| let-7a | 143.5 |
| 5-FU-let-7a | 22.8 |
| miR-145 | 151.8 |
| 5-FU-145 | 11.0 |
| 5-FU | 4612.0 |

**Figure 4.**
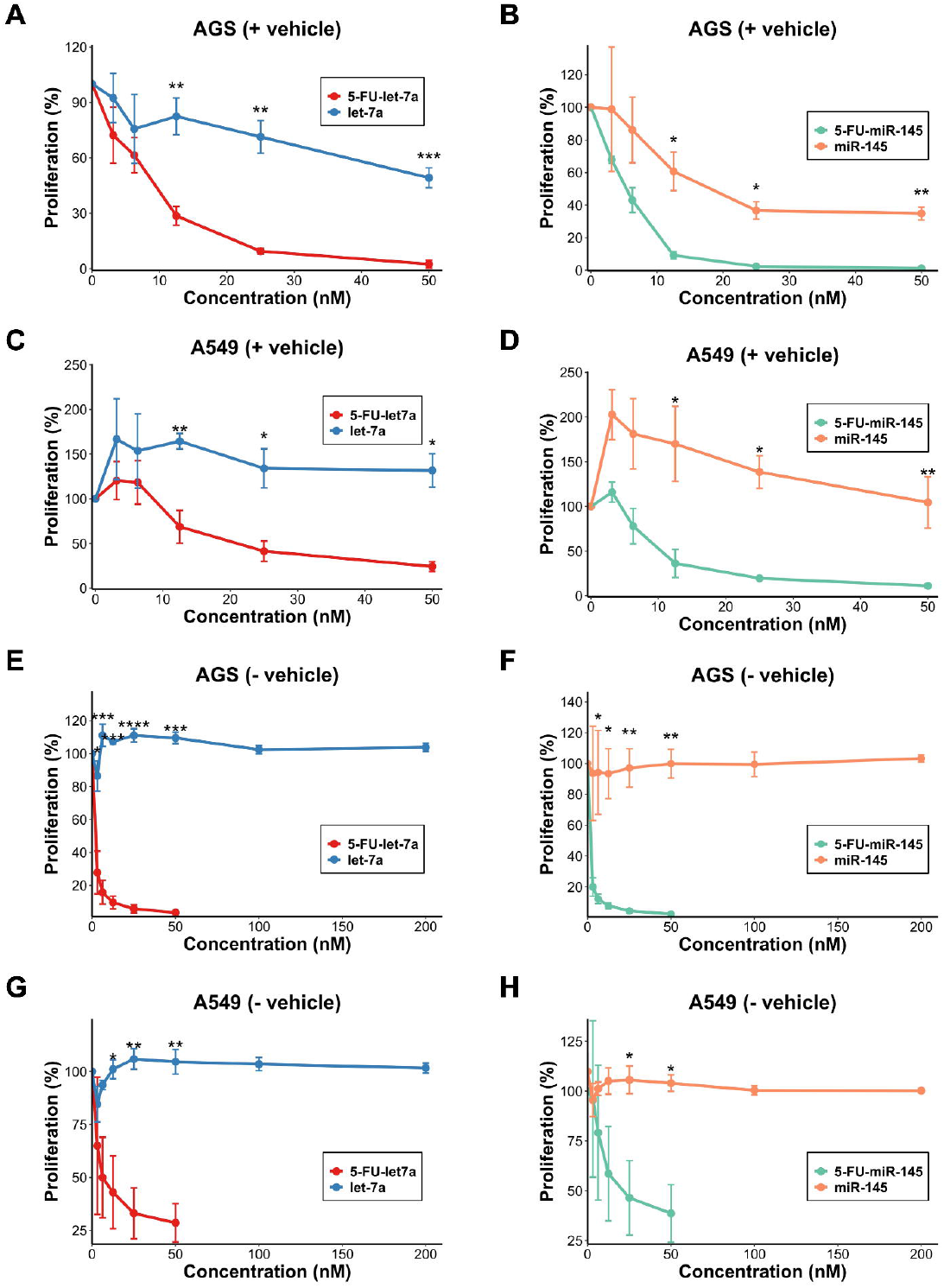
5-FU-modification enhances the tumor suppressor effect of miRNAs and 5-FU-modified miRNA mimetics can enter cells without delivery vehicle. 5-FU-modified miR-145 (5-FU-miR-145) and 5-FU-modified let-7a (5-FU-let7a) inhibits cell proliferation both with and without delivery vehicle in AGS gastric cancer cell line **(A-D)** and in A549 non-small cell lung cancer cell line **(E-H)**. Notably, 5-FU-modified miRNA mimetics appear to be able enter the cell and inhibit cell growth, while unmodified miR-145 and let-7a are only able to inhibit tumor growth in the presence of delivery vehicle. Data are presented as mean ± standard error of the mean and analyzed by Student’s t-test (n = 3).

To further confirm that miRNAs 5-FU-let-7a and 5-FU-miR-145 can enter the cell without the use of transfection vehicle, AGS and A549 cells were treated let-7a and miR-145 without the use of vehicle. There is no inhibition of cell proliferation in cells treated with let-7a and miR-145 without the use of vehicle **(Figure 4E-4H)** compared to cells treated with let-7a and miR-145 in the presence of vehicle (**Figure 4A-4D)**. Notably, 5-FU-let-7a and 5-FU-miR-145 has a significant inhibition of proliferation despite the absence of delivery vehicle **(Figure 4E-4H)**.

The main mechanism of action of 5-FU is forming an irreversible, covalent ternary complex with thymidylate synthase (FdUMP-TS) and tetrahydrofolate. To investigate whether the enhanced efficacy of 5-FU-modified miRNA mimetics are due to the release of 5-FU, western immunoblot analysis of thymidylate synthase (TS) was used to detect the presence of the FdUMP-TS complex. We probed for TS to evaluate whether the 5-FU-modified miRNA mimetics exerted a 5-FU effect. Upon western immunoblot analysis, we observed that 5-FU-let-7a and 5-FU-miR-145 exerts a 5-FU effect as observed by the formation of FdUMP-TS complex, represented as the upper band, in both AGS cells **(Figure 5A & 5B)** and in A549 cells **(Figure 5C & 5D)**.

**Figure 5.**
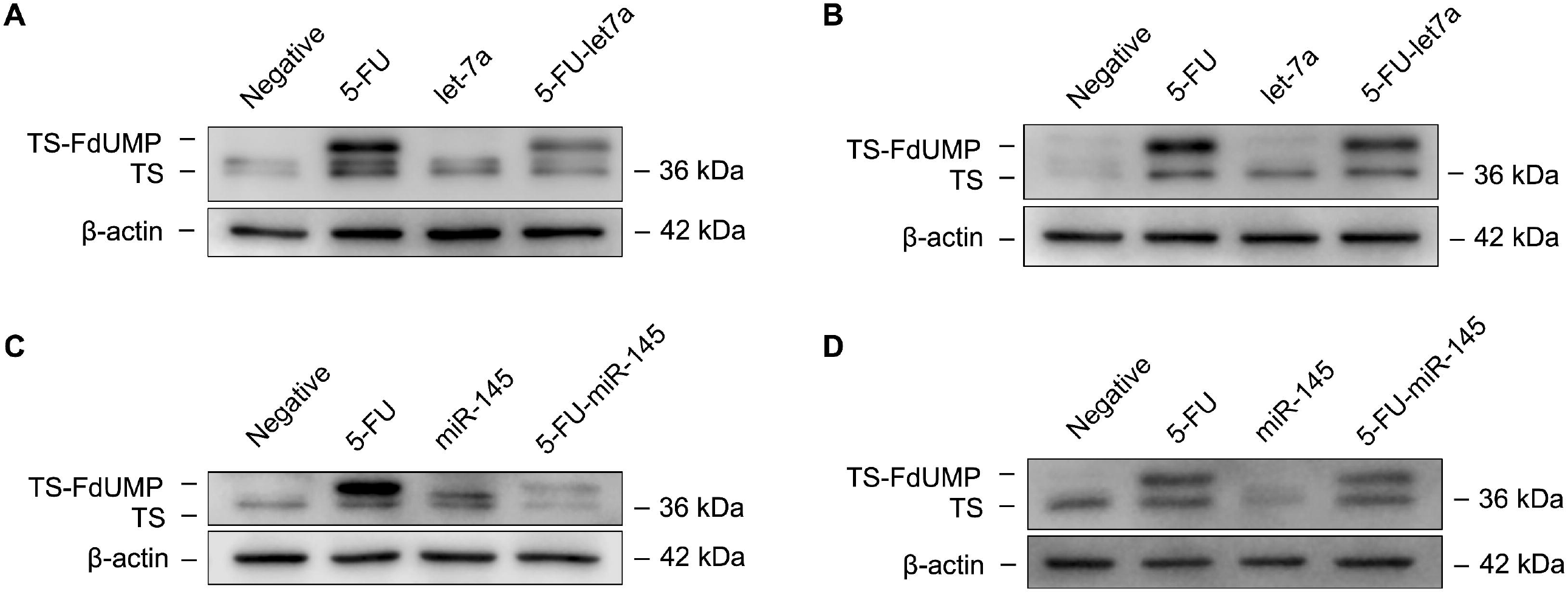
5-FU-let-7a and 5-FU-mi5-145 exhibits 5-FU activity. **(A)** 5-FU-let-7a and **(B)** 5-FU-miR-145 forms a TS-FdUMP ternary complex in AGS gastric cancer cells and A549 lung cancer cells as seen by a band shift upon Western blot of TS. All cells were treated with 50nM of miRNA and the 1µM of 5-FU in AGS cells and 3µM of 5-FU in A549 cells.

### Target specificity of 5-FU-modified miRNA mimetics

To confirm the target specificity of the 5-FU-modified miRNA mimetics, we performed western blot analysis on known targets of let-7a and miR-145. Cyclin-dependent kinase 6 (CDK6) is a direct target of let-7a.^39,40^ Similarly, specificity protein 1 (SP1) is a direct target of miR-145.^41,42^ In both AGS gastric cancer cells and A549 lung cancer cells 5-FU-let-7a decreases CDK6 expression **(Figure 6A & 6B)** and 5-FU-miR-145 decreases SP1 expression under vehicle-free conditions **(Figure 6C & 6D)**. To further confirm that 5-FU-modified miRNA mimetics are effective under vehicle-free conditions, we performed western blot analysis of checkpoint kinase 1 (CHK1) and WEE1 G2 checkpoint kinase (WEE1), two direct targets of miR-15a.^29^ 5-FU-miR-15a decreases the expression of both CHK1 and WEE1, whereas neither miR-15a, 5-FU, nor the combination of the two decreases target protein expression under vehicle-free conditions **(Figure 6E)**.

**Figure 6.**
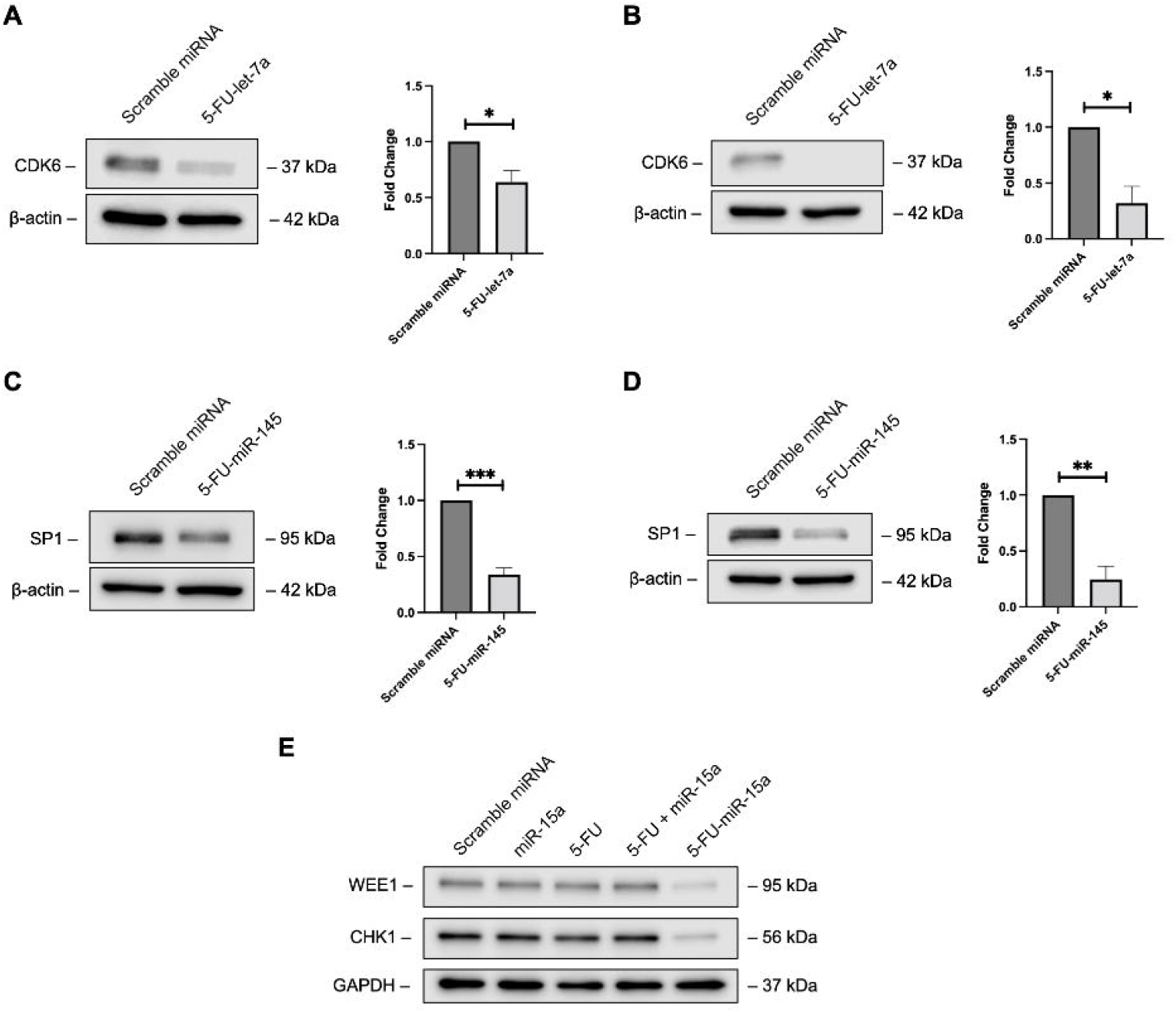
5-FU-modified miRNA mimetics retain target specificity. CDK6 is a known target of let-7a and 5-FU-let-7a can knockdown CDK6 expression in both **(A)** AGS gastric cancer cells and **(B)** A549 lung cancer cells. SP1 is a known target of miR-145 and 5-FU-miR-145 knocks down SP1 expression in both **(C)** AGS gastric cancer cells and **(D)** A549 lung cancer cells under vehicle-free conditions. All cells were treated with 50 nM of either control scramble miRNA or the respective miRNA. **(E)** 5-FU-miR-15a knocks down expression of its targets, CHK1 and WEE1, under vehicle free conditions, whereas miR-15a, 5-FU, or the combination of the two does not in HCT116 colon cancer cells. Data are presented as mean ± standard error of the mean and analyzed by Student’s t-test (n = 3). \**p* < 0.05, \*\**p* < 0.01, \*\*\**p* < 0.001.

### Therapeutic efficacy and safety of 5-FU-modified miRNA mimetics *in vivo*

To demonstrate the therapeutic efficacy of 5-FU modified miRNA mimetics *in vivo*, with and without the delivery vehicle, we choose 5-FU-miR-15a as our top candidate based on the *in vitro* efficacy screening. The *in vivo* effects of a 5-FU-modified miRNA mimetic, 5-FU-miR-15a, was evaluated in a CT-26 syngeneic mouse colon cancer model. Mice were treated with PEI vehicle alone, 40 µg of 5-FU-miR-15a with PEI vehicle, or 40 µg of 5-FU-miR-15a without vehicle. Based on histopathological analysis of the tumors in the mouse lungs, compared to control mice, 5-FU-miR-15a inhibits tumor growth by 58.2% without vehicle (*p* = 0.0066) **(Figure 7B)** and by 97.2% **(Figure 7C)** with vehicle (*p* = < 0.0001). This tumor inhibition was enhanced in the presence of vehicle (*p* = 0.0179) **(Figure 7D)**. To assess whether these animals exhibited any acute toxicity associated with the treatment, their body mass was measured daily during the treatment period. The body mass of the animals stayed within normal healthy limits (<10% change in mass) throughout the treatment period **(Figure 7E)** and liver chemistries did not differ between the treatment groups (**Figure 1**).

**Figure 7.**
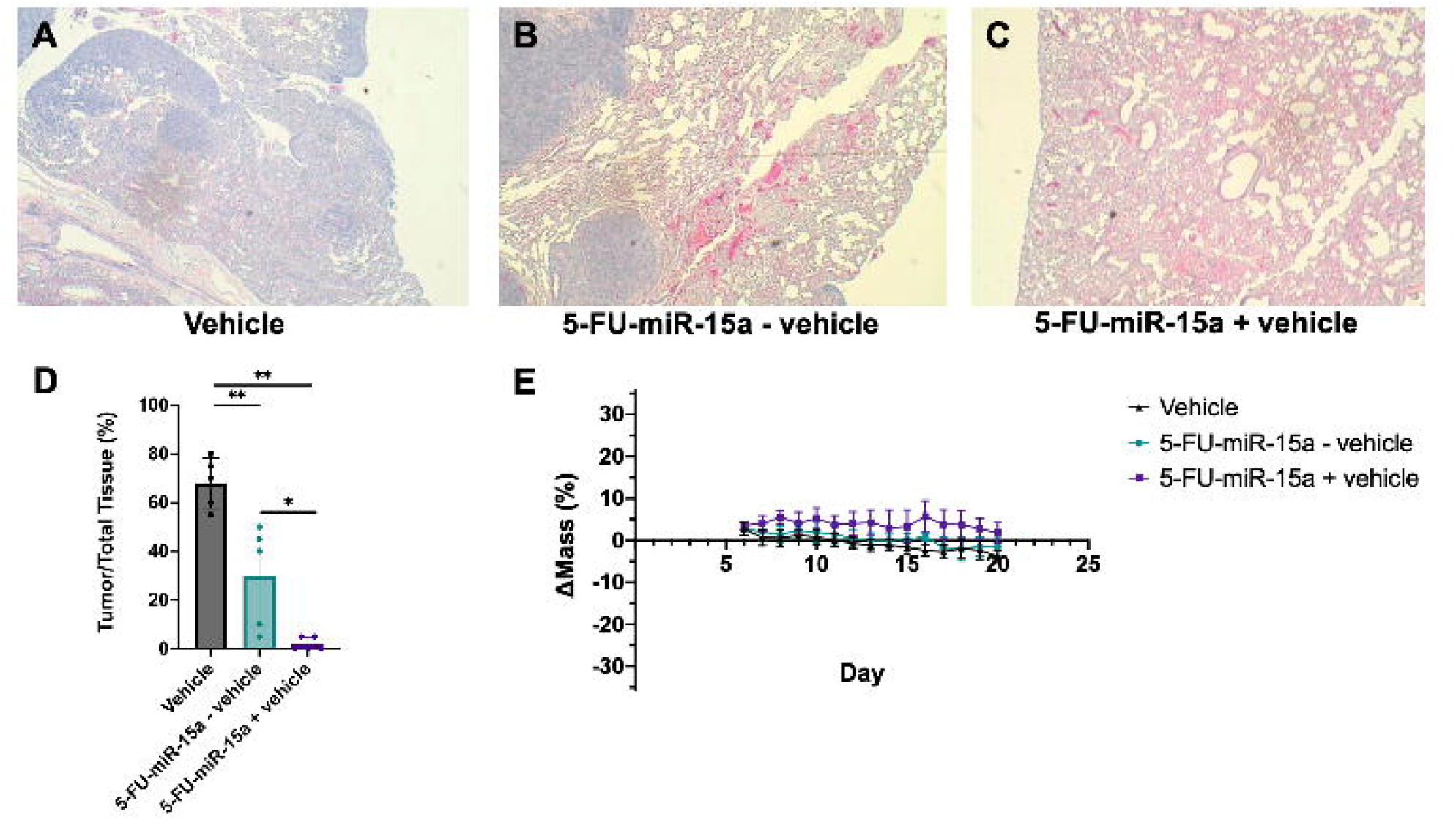
5-FU-miR-15a inhibits tumor growth in an *in vivo syngeneic* colorectal cancer mouse model with and without delivery vehicle. CT-26 tumor allografts established in 8-week-old female BALB/cJ mice via tail vein injection following treatment (40 µg q.o.d.,8 doses). **(a-c)** Representative histological sections of tumors harvested after 20 days and stained with H&E. (**d)** 5-FU-miR-15a inhibits tumor growth with and without PEI, though the efficacy is improved with delivery vehicle (*p* = 0.0179). **(e)** Body weight change was measured as an indicator acute toxicity and no toxicity was observed (< 10% weight loss). Data are presented as mean ± standard deviation and analyzed by Student’s t-test (n = 5). \**p* < 0.05, \*\**p* < 0.01, \*\*\*\**p* < 0.0001.

## Discussion

In this study, we aimed to design a strategy to develop effective miRNA-based therapeutics with enhanced efficacy and stability that can potentially subvert the use of delivery vehicle. One of the key barriers in the development of nucleic acid-based therapeutics is finding methods to effectively introduce these molecules into cells in a minimally cytotoxic manner. While there have been some recent successes with using lipid-based nanoparticles as a vehicle to deliver nucleic-acids in some diseases, toxicities associated with systemic delivery, including immunotoxicity, still exist.^43-47^ The toxicity of delivery vehicle becomes more profound and prohibitory during dose escalation studies as increasing amount of vehicle are needed to deliver the respectively increasing nucleic-acid drug candidates. While delivery vehicles may be relatively safe at low concentrations, this approach relies on an extremely potent biological response to the nucleic acid drug candidate. This is the case in mRNA vaccines, where a small amount of mRNA, and thereby its vehicle, is required for an immunological response. However, in the situation where higher dose of vehicle are needed to pack proportionally larger amounts of nucleic acids, toxicities will become a major bottleneck for safe and effective therapy. Among the 14 currently FDA approved oligonucleotide-based therapeutics, only 6 are delivered systemically. Notably, the use of patisiran (Onpattro) requires premedication consisting of a corticosteroid, acetaminophen, a H1 blocker, and a H2 blocker to prevent infusion-elated reactions.^25^ In this study, we demonstrated an example of toxicities associated with lipid-based nanoparticles, exhibiting a dose-dependent effect on apoptosis **(Figure 2)**.

As a result of these observed toxicities, we sought to enable the use of miRNA-based therapeutics that may be able to get away from the use of large amounts of delivery vehicle or eliminating their use altogether. In this study, we focused our efforts to demonstrate the broad potential of miRNA-based therapeutics by enhancing its deliverability. This novel approach was approached by modified guide strand of tumor suppressor miRNA by replacing the uracils with 5-FUs (**Figure 1**). This approach has minimal alterations of the miRNA molecule, as the single hydrogen to fluorine substitution at the 5-position of uracil does not interfere with its Watson-Crick base pairing with adenine. By incorporation of 5-FU to miRNA, we were able to deliver the 5-FU modified miRNA mimetics under without use of delivery vehicle *in vitro* **(Figure 3)**. These modifications also enhanced efficacy of native miRNA by combine the therapeutic effects of miRNA and 5-FU into one entity, as well as creating a more stable molecule against degradation.^30^ We have previously demonstrated in colon cancer, pancreatic, and breast cancer that 5-FU modification of miRNAs have several key features such as retaining target specificity, vehicle-free delivery, enhanced potency, and increased intracellular stability.^28,30,31,48^ To demonstrate the general applicability of this approach to other tumor suppressor miRNAs, we chose 12 well-studied tumor suppressor miRNAs and applied our 5-FU modification strategy. We screened these 12 different 5-FU-modified miRNA mimetics in several other major cancer types including breast cancer, gastric cancer, leukemia, lung cancer, and pancreatic cancer. While it is impractical to screen all potential tumor suppressor miRNAs, the tumor suppressor miRNAs selected in this study have been previously shown to inhibit cancer growth, disease progression, metastasis, and/or drug resistance.^38^ Their biological roles and their targets are also defined, well-investigated, and briefly summarized below. As previously stated, miR-15 regulates apoptosis by targeting BCL2. Similarly, miR-129 can also regulate BCL2 expression and other protumorigenic proteins such as high-motility group box-1 (HMGB-1) and CDK6.^7,49,50^ The let-7 family regulates the RAS oncogene expression and is found to be downregulated in multiple cancer types.^6,51,52^ miR-34 directly targets P53 and thus it has an impact on multiple pathways including tumor proliferation, apoptosis, and cell cycle. ^35,53-57^ miR-140 regulates stemness through some of its targets HDAC4, SOX9, and ALDH1. miR-145 expression is found to be highly downregulated in colon cancer^9^ and regulates cancer growth, in several tumor types, through its targets that include IGF1R, MYO6, and SP1.^9,41,58-62^ miR-194 inhibits metastasis and invasion by inhibiting BMP1, p27^kip1^, and RBX1.^63-65^ The miR-200 family (including miR-200a/b/c) has been shown to inhibit EMT by downregulating ZEB1 and ZEB2 in multiple tumor types including breast, gastric, lung, and pancreatic cancer.^8,36,66-69^ miR-215 is a cell cycle regulator and delivering miR-215 to cancer cells causes cell cycle arrest in several cancer types.^11,70,71^ Lastly, miR-506 also regulates cancer progression and invasion by targeting the NF-κB pathway, SNAI1, and YAP1.^33,34,72,73^ In summary, these tumor suppressor miRNAs have diverse and sometimes context-dependent biological functions thus it is important to screen multiple therapeutic candidates.

During this screen, we observed that all 12 5-FU-modified miRNA mimetics were able to inhibit cancer cell proliferation at a concentration of 50 nM without use of delivery vehicle (**Figure 3**). Although we noticed that each unique 5-FU modified miRNA mimetic has different levels of efficacy *in vitro*, interestingly, certain cell lines—AGS gastric cancer cells and REH leukemia cells—were highly sensitive to all 5-FU-modified-miRNA-mimetic treatment **(Figure 3A &3D)**. This raises an interesting finding that for gastric cancer and leukemia, 5-FU-modified miRNA mimetics are all strong potential therapeutic candidates despite the different targets and pathways that are impacted by each miRNA. Although we have demonstrated previously that 5-FU modification of miRNAs (miR-129, miR-15a, miR-489) does not alter target specificity^28-31^, to further validate that target specificity is maintained with this approach, we selected two additional miRNAs to investigate further—let-7a and miR-145—as they are previously reported to be markers of aggression and prognosticators in gastric cancer^41,42,60,74-76^ and in lung cancer^58,59,61,77,78^. Our results demonstrate that 5-FU-let-7a and 5-FU-miR-145 retain target specificity to CDK6 and SP1 respectively, indicating that there is successful knockdown of their previously reported target genes **(Figure 6A-6D)**. To further validate that 5-FU-modified miRNA mimetics retain target specificity and can do so under vehicle-free conditions, we investigated two direct targets of miR-15a, CHK1 and WEE1. Our results demonstrated that 5-FU-miR-15a can knock down target expression, without delivery vehicle, while neither miR-15a, 5-FU, nor the combination of two are able to knock down expression **(Figure 6E)**. Collectively, our results demonstrated that 5-FU modification of tumor suppressor miRNAs retain target specificity. We also observe that 5-FU-let-7a and 5-FU-miR-145 form a FdUMP-TS complex, demonstrating that 5-FU is indeed released, potentially as a breakdown product of these mimetics, and can exert 5-FU activity in cells **(Figure 5)**. Taken together, our results show that 5-FU-modified miRNA mimetics are potent therapeutic molecules, effective in the nanomolar range and that they are more effective at inhibiting cancer cell proliferation than unmodified miRNAs.

To demonstrate the therapeutic potential of 5-FU modified miRNA mimetics *in vivo*, we selected 5-FU-miR-15a based on our *in vitro* screening using an immunocompetent syngenetic mouse colon cancer model. Using a tail vein injection syngenetic colon cancer mouse model, we observed that 5-FU-miR-15a can inhibit cancer cell growth and lung metastasis both with and without delivery vehicle **(Figure 7A-7D)**. This model was selected as a proof-of-concept to model efficacy in a metastatic disease setting, the largest burden on cancer morbidity and mortality. It is worth noting that the therapeutic efficacy of 5-FU-miR-15a can be further enhanced with a low concentration of a polyethyleneimine LNP delivery vehicle. This is an expected outcome of this investigation, as we sought to create miRNA-based therapeutics that relies only on a little to no delivery vehicle, key to avoiding potential toxicities **(Figure 7)**. This is the first time, to our best knowledge, miRNA-based cancer therapeutics have been demonstrated to be effective without the aid of delivery vehicle *in vivo*. Mice treated with 5-FU-miR-15a with and without delivery vehicle show no significant weight and hair loss **(Figure 7E)**. Liver chemistries also did not differ between treatment groups, potentially representing a lack of hepatotoxicity from 5-FU-modified miRNA mimetic treatment **(Figure S1)**. Additionally, there were also no behavioral changes, such as loss of appetite, among the treated mice. The demonstration of efficacy in the presence and absence of delivery vehicle may allow for flexibility in the optimization of the formulation of the mimetics for future clinical use, and to avoid the bottleneck of toxicity. Future therapeutic development can consider the reduction or elimination of vehicle thus potentially avoiding side effects due to toxicity as seen in systemic chemotherapeutic regimens. Similarly, low toxicities may give rise to a larger therapeutic window of 5-FU-modified miRNA mimetics and further work must be completed to optimize the dosage of these mimetics.

Previous studies from our group have compared a few different modification strategies of miRNAs, including varying the number of 5-FU substitutions and locations of the moiety and have demonstrated that the substation of all uracil bases on the guide strand appear to be the most effective.^30^ A potential mechanism of action of the observed enhanced efficacy and deliverability of the 5-FU-modified miRNA mimetics is due to the increased lipophilicity that is conferred with the addition of fluorine to drug candidates, which is a strategy that has been used for improving lipophilicity of small molecule compounds.^79,80^ This increase in lipophilicity may allow for the miRNA—negatively charged and typically unable to cross the cell membrane—to cross the cell membrane. Our approach takes advantage of 5-FU as an active anti-cancer therapeutic compound and the fluorine group will also enhance the deliverability of a nucleic-acid based miRNA tumor suppressor. There are various modification strategies in nucleic acid drug development—especially in antisense oligonucleotides—including 2’-*O*-methyl that confers more target affinity and 2’-fluoro that confers more nuclease resistance.^81,82^ We have previously observed that 5-FU modification in miR-129 confers additional intracellular stability^30^, however in this study, we did not observe an increase in half-life of 5-FU modified miRNA mimetics in cell culture media supplemented with 10% FBS (data not shown). Notably, we did not make any additional modifications to attempt to preserve the native miRNA tumor suppressor function. Future studies can consider some of these approaches to optimize the design of these drug candidates. It is also difficult to tease apart the individual contributions to the anti-tumor phenotype of the 5-FU and the tumor suppressor miRNAs that make up the 5-FU-modified miRNA mimetics. Future studies evaluating 5-FU-modified anti-miRNAs may help begin answering some of those mechanisms. Similarly, while the 5-FU-modified miRNA mimetics appear to be broadly effective, the selection of specific 5-FU-modified miRNA mimetics for further pre-clinical development should consider miRNA targets of oncogenic signaling pathways and the specific cancer type.

In summary, our study demonstrates that 5-FU-modified miRNA mimetics display broad therapeutic potential, as they are effective in several different cancer types without the aid of transfection vehicle. This study expands on previous work on 5-FU-modified miRNA mimetics, showing that they are efficacious in additional cancer types including gastric cancer, lung cancer, and leukemia. Notably, 5-FU-modified miRNA mimetics retain their mRNA target specificity and appears to be well tolerated in our animal studies. Overall, 5-FU modification of miRNAs may serve as a novel technology platform for broad miRNA-based therapeutic development.

## Materials and Methods

### Design and synthesis of the 5-FU-modified miRNA mimetics

The 5-FU-modified miRNA mimetics were designed and synthesized by substituting uracil with 5-fluorouracil (5-FU) on the guide strand of the miRNA. The passenger strand was left unmodified to avoid any potential off-target effects and to preserve miRNA function. Oligonucleotides with these modifications as well as their corresponding passenger strand were purchased from Dharmacon (Horizon Discovery). Both strands of oligonucleotides were HPLC purified. The guide strands and passenger strands were then annealed prior to use.

### Cell culture

All cell lines were obtained from ATCC and are derived from human cells. AGS gastric cancer and A549 lung cancer cells were cultured in Ham’s F-12K (Kaighn’s) Medium supplemented with 10% fetal bovine serum (FBS). SKBR3 breast cancer cells and HCT116 colon cancer cells were cultured in McCoy’s 5A Medium supplemented with 10% FBS. REH leukemia cells and AsPC-1 pancreatic cancer cells were cultured in RPMI-1640 Medium supplemented with 10% FBS. Hs766T and PANC-1 pancreatic cancer cells were cultured in Dulbecco’s Modified Eagle Medium (DMEM) supplemented with 10% FBS.

### Western immunoblot analysis

AGS, A549, and HCT116 cells were seeded onto 6-well plates at a cell density of 100,000 cells per well. 24 hours later, the cells were transfected with 50 nM of miRNA under vehicle-free conditions. HCT116 cells were also treated with a 50 nM miR-15a control condiditon, a 350 nM of 5-FU (equivalent 5-FU concentration in the 5-FU modified miRNA), and the combination of the two. 72 hours later, cells were lysed with RIPA buffer and the protein samples were used for western immunoblotting. Proteins were probed with anti-thymidylate synthase antibody (Millipore, Cat:MAB4130, 1:500), anti-CDK6 antibody (Cell Signalling, Cat:13331, 1:10,000), anti-SP1 antibody (Abcam, Cat:ab124804, 1:10,000), β-actin antibody (Invitrogen, Cat:A5441, 1:10,000,000), anti-CHK1 antibody (Cell Signalling, Cat:, 1:1000), anti-WEE1 antibody (Cell Signalling, Cat:, 1:1000), and anti-GAPDH antibody (Santa Cruz, Cat:, 1:100,000). Protein bands were visualized using a LI-COR Biosciences Odyssey FC imaging system after the addition of SuperSignal West Pico chemiluminescent substrate (Thermo Fisher Scientific). Proteins were quantified with Image Studio Version 5.2.4 (LI-COR Biosciences). For the 5-FU condition of the thymidylate synthase blot, AGS cells were treated with 1 µM of 5-FU and A549 cells were treated with 3 µM of 5-FU.

### Apoptosis assay

AGS and A549 cells were treated with a polyethyleneimine-based lipid nanoparticle (*in vivo*-jetPEI®, Polyplus). 48 hours later, the cells were stained with Annexin V (Thermo Fisher Scientific) and apoptotic cells were quantified by flow cytometric analysis.

### Cell proliferation assay

For vehicle-free miRNA treatment, cells were seeded onto 96 well plates at 1000 cells per well. 24 hours after seeding, 5-FU-modified miRNA mimetics were added to the cells. These cells were incubated for 24 hours and then the media was changed to fresh media supplemented with 10% dialzed FBS. For miRNA treatement with vehicle, cells were seeded onto 6 well plates at 100,000 cells per well. MiRNAs were combined with Oligofectamine™ (Thermo Fisher Scientific) and then added to the cells. 24 hours later, the cells were trypsized and re-seeded onto a 96 well plate at 1000 cells per well. Cell viability was measured 6 days post transfection using WST-1 reagent (Roche). Cells were incubated with 10 μl of WST-1 per 100 μl of media for 1 hour and absorbance was read at 450 and 630 nm. The O.D. was calculated by subtracting the absorbance at 630 nm from that at 450 nm and the relative proliferation was calculated by normalizing the O.D. to negative control.

### Syngeneic mouse model

8-week-old female BALB/cJ mice (Jackson Labs 000651) were inoculated with 5 × 10^5^ CT-26 syngeneic colon cancer cells suspended in 0.1 mL of PBS via tail vein injection. Five days post inoculation, mice were treated with either 9.6 µM of vehicle alone (*in vivo*-jetPEI®, Polyplus), 5-FU-miR-15a miRNA (40 µg) with vehicle, or 5-FU-miR-15a (40 µg) without vehicle on alternating days for a total of 8 doses, with 5 mice per treatment group. Vehicle concentrations were selected in a non-toxic range as per manufacturer recommendations. All treatments were diluted in 5% glucose to a final volume of 0.1 mL and given via tail vein injection. Mouse tumors were harvested from the lungs at the day 20 endpoint of the study and tumors were formalin fixed, paraffin embedded, and mounted onto slides for staining with H&E. Slides were assessed by a board certified pathologist and scored for % tumor content. Blood was also harvested at the endpoint and was sent to a clinical chemistry lab for subsequent liver enzyme analysis.

### Statistical analysis

The quantitative data were presented as mean value ± standard error of the mean of at least 3 independent experiments in all in vitro studies. Data were analyzed by two-tailed Student’s t-test. The results of the animal studies presented as mean value ± standard deviation. A p value of less than 0.05 were considered to be statistically significant (\**p* < 0.05, \*\**p* < 0.01, \*\*\**p* < 0.001, \*\*\*\**p* < 0.0001).

## Acknowledgements

We would like to thank Matthew Godwin and Zachary Ye for their support in this project. This study was supported by NIH/NCI R01CA197098-01 (J. Ju), Curamir Therapeutics Inc. (J. Ju), and VA Merit Award BX005260-01 (J. Ju).

## Author Contributions

Conceptualization, J.G.Y., A.F., and J.J; Methodology, J.G.Y. and J.J; Investigation, J.G.Y, A.F., G.H; Writing – Original Draft, J.G.Y. and J.J.; Writing Reviewing & Editing, J.G.Y., A.F., G.H., and J.J.; Supervision J.J., Project Administration, L.C., J.J.; Funding Acquisition, J.J.

