## Supplemental Figure 1 for "Development of 5-FU-modified tumor suppressor microRNAs as a platform for novel microRNA-based cancer therapeutics"

**Supplemental Information**

**Supplemental Figure 1. Liver enzymes levels of *in vivo syngeneic* colorectal cancer mouse model.** ALT, ALP, and AST levels of the mice in the treatment group were sampled from mice sera collected at the study endpoint. All levels did not statistically differ across the treatment groups ($p > 0.05$). Data are presented as mean ± standard error and analyzed by Student's t-test (n = 5).

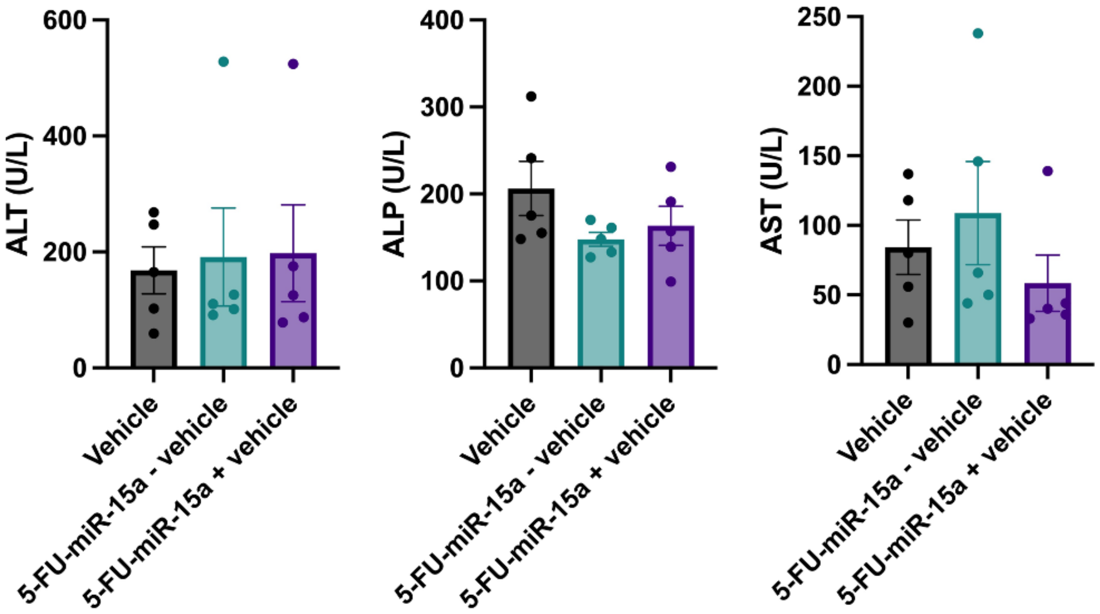
